# A miniaturized *Arabidopsis thaliana* - *Botrytis cinerea* seedling pathosystem and the GreenDayz workflow for longitudinal image-based phenotyping

**DOI:** 10.64898/2026.10.11.758349

**Authors:** Ines Tguafaiti, Christophe Bruel, Mathias Choquer, Caroline Bournaud, Nathalie Poussereau, François-Xavier Gillet

## Abstract

Plant-pathogen interactions are strongly influenced by host developmental stage, yet many *Arabidopsis thaliana-Botrytis cinerea* assays remain centred on mature rosette leaves and rely on inoculum loads that may exceed those involved in a primary infection. We established a miniaturized *in vitro A. thaliana-B. cinerea* seedling pathosystem initiated with a very low conidial dose (nominally ∼25 conidia per plate) compatible with both destructive molecular sampling and repeated, non-destructive RGB imaging. The pathosystem was coupled to a low-cost smartphone photo station and to GreenDayZ Phenotyping, a guided image-analysis application with an intuitive graphical interface that standardizes longitudinal RGB image processing and extracts quantitative colour-based phenotypes. Together, the pathosystem and GreenDayZ Phenotyping provide an accessible and reproducible framework for longitudinal phenotyping of low-inoculum seedling infections, in which image-derived trajectories can be aligned with independent molecular readouts.

## Introduction

Plant-pathogen interactions are strongly influenced by the host developmental stage. Immune competence, hormone signalling and the physiological resources available for defence change throughout plant development, generating age-dependent differences in susceptibility or resistance to pathogens (Huot et al., 2014 ; Develey-Rivière and Galiana, 2007). In *Arabidopsis thaliana*, experimental studies of fungal infection have largely focused on mature rosette leaves, whereas comparatively fewer studies have examined interactions at the seedling stage (e.g. Ortega-Cuadros et al., 2022; Sun et al., 2022). Seedlings nevertheless provide a particularly interesting experimental window: they represent an intact, whole-plant system in which the immune system is still undergoing developmental maturation, while their small size allows host-pathogen interactions to be examined at high spatial and temporal resolution. In addition, infection can develop fastly and simultaneously at multiple sites within the seedling, offering opportunities to investigate heterogeneity in host-pathogen interactions at the whole-plant scale.

The seedling format also provides important experimental advantages for studying the earliest stages of infection. Miniaturization enables large numbers of samples to be handled in parallel and facilitates kinetic designs that would rapidly become impractical with mature plants. At these early stages, the low fungal-to-host biomass ratio can substantially dilute fungal-derived molecules in bulk samples, making fine molecular characterization of the interaction technically challenging. This is particularly relevant for *Botrytis cinerea*, for which early colonization can precede extensive tissue collapse and may involve only a small fungal biomass (Tguafaiti et al, unpublished). Molecular approaches such as qPCR, phytohormone profiling or dual RNA sequencing are therefore confronted with a technical trade-off: increasing inoculum facilitates fungal detection but can shift the experiment towards infection conditions that are less representative of a primary infection, whereas low inoculum better preserves the early interaction but generates a weak pathogen-derived molecular signal (Girard et al., 2016). A miniaturized seedling pathosystem could provide a means of partially alleviating this constraint by combining low-dose infection, large replicate numbers and the possibility of pooling whole seedlings to obtain sufficient biological material for molecular analyses.

Such kinetic experiments, however, create a second requirement. Relating molecular measurements to disease progression requires a non-destructive phenotypic readout that can be repeatedly acquired from the same experimental units throughout the infection time course. RGB photography is particularly attractive in this context because it is inexpensive, non-invasive and readily scalable. Image-based plant phenotyping has consequently generated a wide range of approaches for measuring projected plant area, greenness, lesion development and other colour-derived traits, ranging from vegetation indices to dedicated plant-image analysis pipelines (Easlon *et al*., 2014; Pavicic *et al*., 2021; Signorelli *et al.,* 2025). More recently, user-oriented tools have facilitated the extraction of phenotypic information from RGB images without requiring advanced computer-vision expertise (Lee *et al.,* 2024; Hu *et al*., 2025). Nevertheless, many available approaches either address a specific phenotype, rely on relatively standardized imaging conditions, or require multiple independent processing steps. There remains a practical need for workflows that can manage complete longitudinal experiments, incorporate acquisition-related quality control and normalization, retain visual verification of segmentation.

To address these combined biological and analytical constraints, we developed a miniaturized sterile *A. thaliana–B. cinerea* seedling pathosystem designed for low-dose inoculation, rapid destructive sampling and repeated longitudinal imaging. Seedlings are grown as grouped populations on removable nylon supports, allowing experimental units to be harvested rapidly for molecular analyses while parallel plates can be followed non-destructively over time. Because such kinetic experiments rapidly generate large image series, we coupled the pathosystem to a standardized low-cost smartphone imaging setup and to GreenDayZ – Phenotyping, a guided image-analysis application designed to manage complete longitudinal datasets, reduce acquisition-related variability and extract reproducible colour-based phenotypes without requiring specialist expertise in image analysis or programming. Together, these components provide a practical framework for aligning longitudinal phenotypic trajectories with independent molecular measurements during low-inoculum seedling infection.

## Results

### A miniaturized in vitro seedling pathosystem compatible with destructive sampling and repeated imaging

To investigate the progression of the A. thaliana–B. cinerea interaction over several days, we established a miniaturized in vitro seedling pathosystem based on grouped seedling populations grown on 100-µm nylon mesh in 55-mm Petri dishes (Fig. 1; Supplementary Fig. S1). The nylon support allows the entire seedling population to be rapidly harvested with limited agar carry-over and mechanical disturbance. This rapid sampling is particularly important for molecular analyses, as it minimizes the interval between removal from growth conditions and sample freezing and thereby limits handling-induced physiological or transcriptional responses in both host and pathogen. After seedling establishment, plates were inoculated centrally with a single 10-µL drop delivering approximately 25 B. cinerea conidia per plate. Because conidia sediment rapidly at such low concentrations, the inoculum was maintained under continuous stirring and the effective delivered dose was independently controlled for each inoculation session.

**Figure 1.**
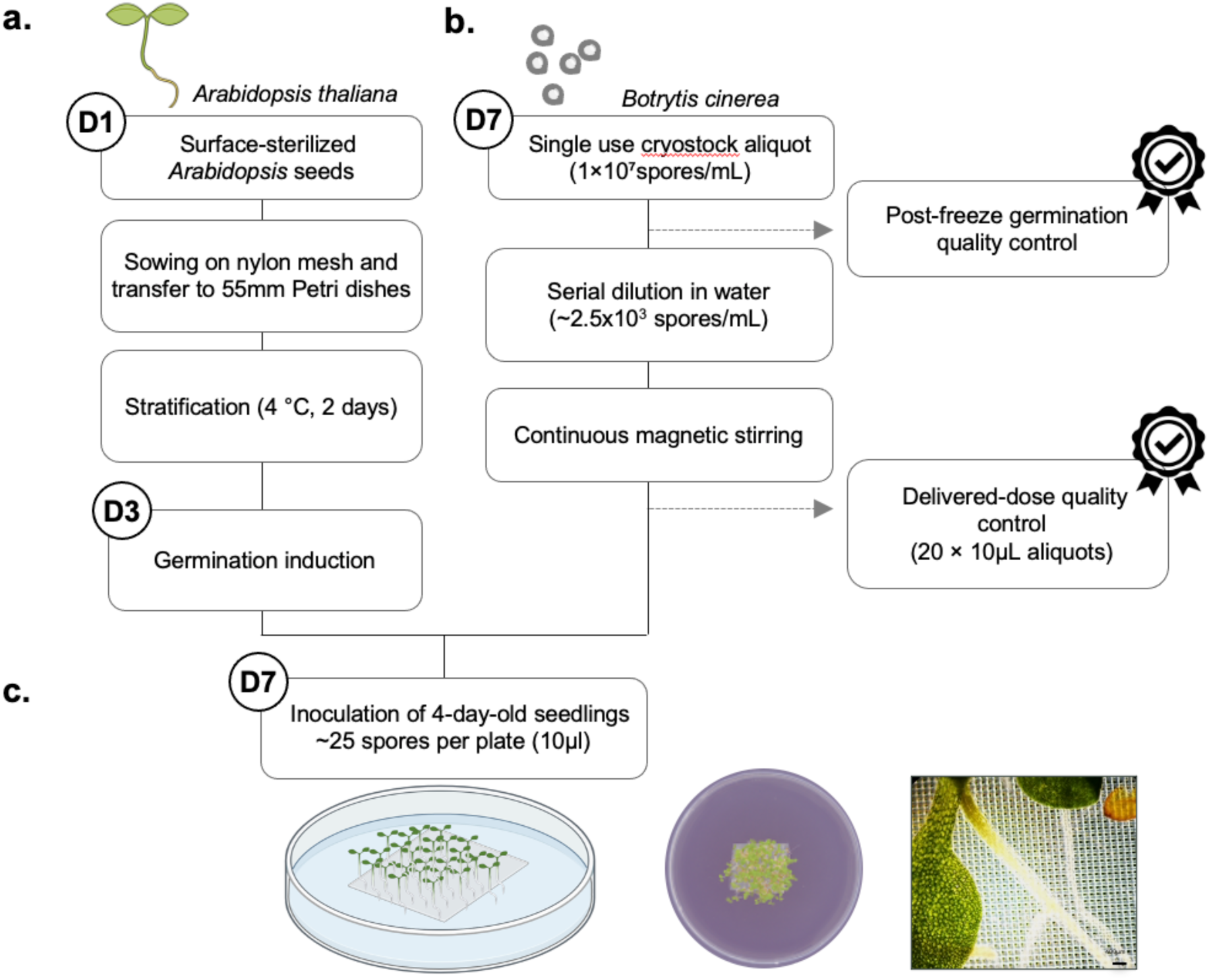
Miniaturized low-inoculum *Arabidopsis thaliana-Botrytis cinerea* seedling pathosystem. **(a)** Establishment of grouped *A. thaliana* seedlings on 100µm nylon mesh and transfer to 55mm Petri dishes containing nutrient agar medium. The nylon support facilitates rapid recovery of the complete seedling population for downstream molecular analyses. **(b)** Preparation of a low-dose *B. cinerea* inoculum from a single-use frozen conidial stock. The working suspension is serially diluted to approximately 2.5 × 10³ conidia mL⁻¹ and maintained under continuous magnetic stirring to limit conidial sedimentation. Inoculum quality is controlled by post-freeze germination assessment and by independent counting of 20 delivered 10µL aliquots. **(c)** Four-day-old seedling populations are inoculated with a single central 10µL drop delivering approximately 25 conidia per plate. Mock controls receive sterile water. Detailed seed-sowing and transfer procedures are shown in Supplementary Fig. S1.

This miniaturized format enables large kinetic experiments to be conducted in parallel. For example, a destructive time-course experiment comparing infected and non-infected seedlings over seven days with six biological replicates per condition already requires 84 experimental plates, before adding backup plates to compensate for irregular seed deposition, contamination or other quality-control exclusions. Such experimental designs rapidly generate large longitudinal image series, creating the need for a standardized acquisition setup and a dedicated image-analysis workflow.

### A simple DIY photo station for reproducible Petri-dish imaging

A low-cost photo station was assembled directly inside the sterile work area used for plate handling (Fig. 2). The setup combines a fixed smartphone position, a consumer LED ring light and a black-and-white Plexiglas support that reproducibly positions the Petri dish. The black background increases contrast between plant material and the surrounding agar/background, thereby facilitating subsequent image segmentation, whereas the surrounding white area provides a neutral reference for white-balance normalization. A small colour chart integrated into the support provides additional reference colours for calibration and technical validation. The photo station constrains the main geometric and illumination parameters and embeds reference information within each image, allowing residual colour variation to be quantified and corrected downstream.

**Figure 2.**
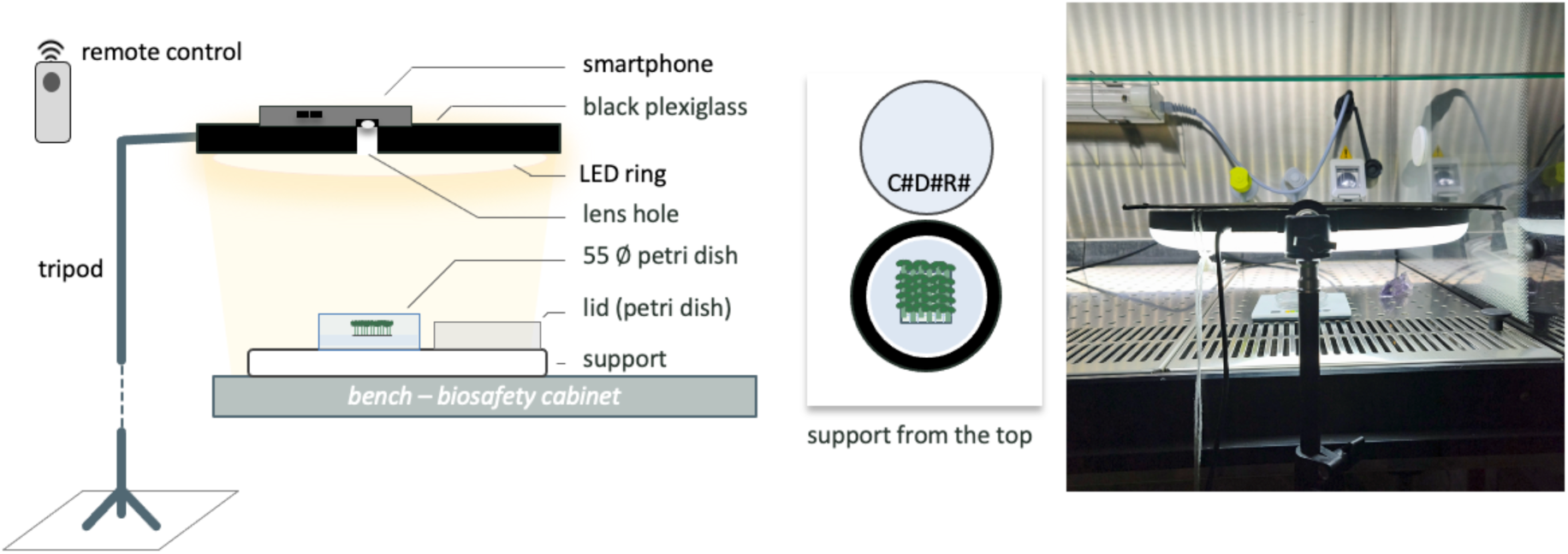
DIY smartphone imaging station for standardized Petri-dish acquisition inside a microbiological safety cabinet. A smartphone is positioned above a horizontally oriented LED ring light using a fixed black Plexiglas support, with the camera aligned through a dedicated lens opening. The 55mm Petri dish is positioned on a black background within a white reference area that also includes a colour chart. This arrangement standardizes plate positioning and illumination while retaining reference regions for downstream image normalization and technical validation. Each Petri-dish lid carries a sample code visible in the acquired image (e.g. C1D1R1, indicating condition 1, day post inoculation 1 and biological replicate 1), thereby preserving sample identity directly within the photographic record. Image filenames additionally include the acquisition date and time (YYYYMMDD_HHMMSS), providing an independent timestamp and strengthening traceability across longitudinal image series. The Petri-dish lid is briefly removed during acquisition to minimize reflections from transparent plastic, and a Bluetooth shutter allows rapid image capture without touching the smartphone or support.

### GreenDayZ-Phenotyping provides a guided image-analysis workflow

GreenDayZ – Phenotyping combines four broad functions: dataset quality control, image preprocessing, image processing and feature extraction, and visual and analytical reporting (Fig. 3). The application is implemented as a local Python-based graphical interface designed to run on both macOS and Windows. Its interface mirrors the sequential organization of the analytical workflow, allowing users to move from one processing module to the next while retaining access to step-specific settings, previews, quality-control information and outputs. Each major processing step can be previewed or validated before batch processing continues, while quality-control images, summary tables and analytical outputs are stored with the experimental dataset.

**Figure 3.**
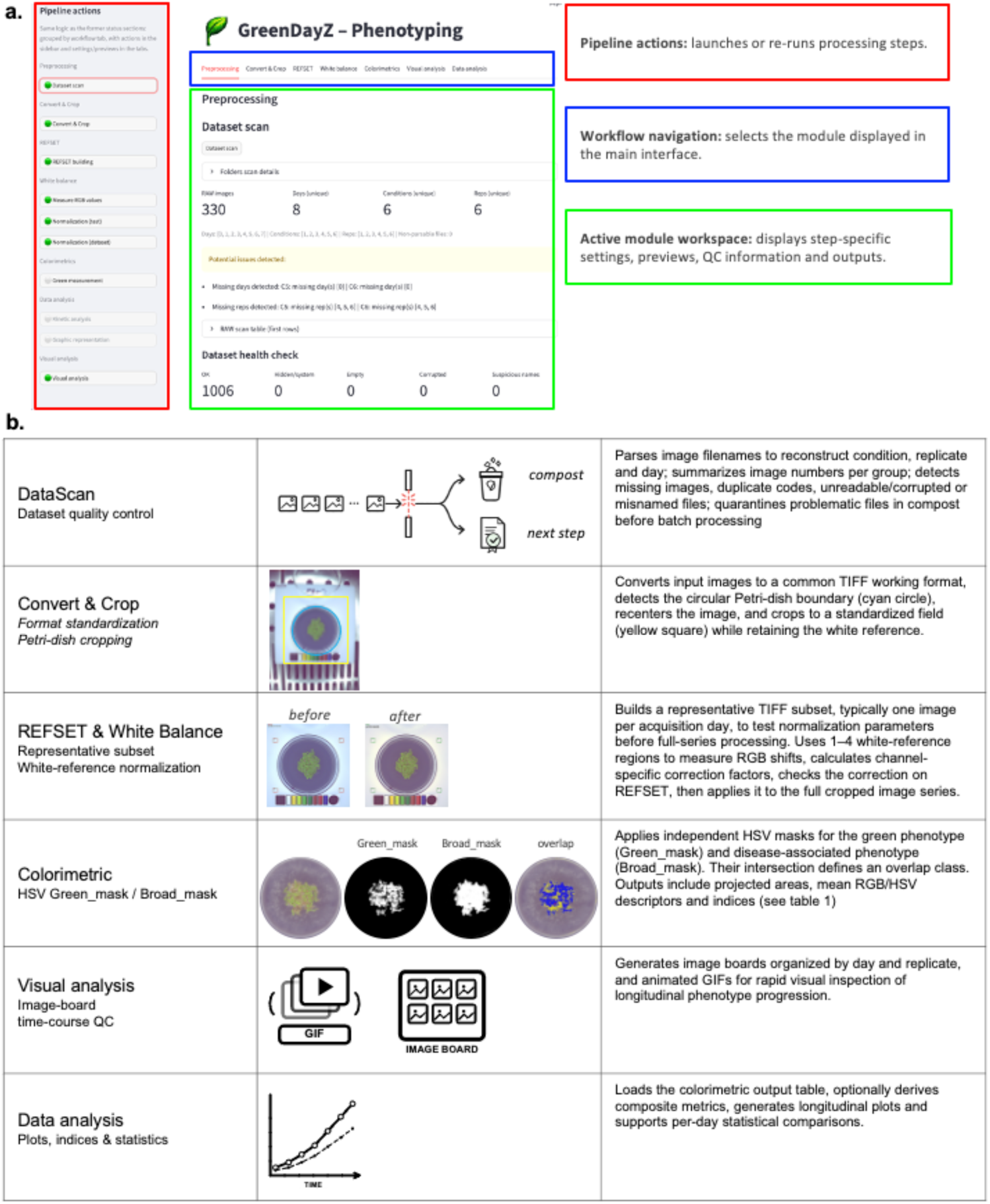
GreenDayZ - Phenotyping image-analysis workflow and representative interface outputs. **(a)** Graphical interface, shown on the dataset-scan step, organized into three areas: pipeline actions (red), workflow navigation (blue) and the active module workspace (green), which displays step-specific settings, previews and quality-control outputs. **(b)** The six sequential modules, each with a representative output: DataScan controls dataset integrity, Convert & Crop standardizes the format and crops each plate to a common field, REFSET & White Balance applies a white-reference normalization, Colorimetric extracts the Green_mask, Broad_mask and their overlap, Visual analysis generates image boards and animated GIFs, and Data analysis produces longitudinal plots and per-day statistics. Colorimetric readouts are detailed in Table 1.

**Table 1.** Colorimetric readouts derived from the GreenDays Colorimetrics workflow. Colorimetrics generates both raw measurements and derived indices for each analysed image and exports them as a tabular dataset in Excel-compatible format. The core readouts retained here summarize complementary aspects of the green phenotype and its transition toward non-green colour classes. A mask selects which pixels are analysed but does not modify their RGB intensities. Definitions used below: Green_mask = mask selecting green-classified pixels; Broad_mask = broad colour-transition mask; ROI = Petri-dish region of interest; Green_area = area classified by the Green_mask; Broad_area = area classified by the Broad_mask; Overlap_area = area shared by the Green_mask and Broad_mask; ROI_area = total analysed ROI area; R, G and B = mean RGB intensities within the Green_mask; H = Hue value of a pixel within the Green_mask.

| readout | Equation | Interpretation |
| --- | --- | --- |
| Green area fraction | $\text{Green\_area} / \text{ROI\_area}$ | **How much green plant surface is visible in the analysed image?**<br>Fraction of ROI pixels belonging to the Green_mask. It reports changes in the projected green plant surface over time. |
| Green-state fraction | $\text{Overlap\_area} / \text{Broad\_area}$ | **Within the Broad_mask, what fraction of the area still overlaps with the Green_mask?** This ratio compares the area shared by the Green_mask and Broad_mask (Overlap_area) with the total area detected by the Broad_mask (Broad_area). A value close to 1 means that most Broad_mask pixels are also classified as green; a decrease indicates that an increasing fraction of the Broad_mask no longer overlaps with the Green_mask. |
| Non-green fraction | $(\text{Broad\_area} - \text{Overlap\_area}) / \text{Broad\_area}$<br>$= 1 - \text{Green-state fraction}$ | **Within the Broad_mask, what fraction is no longer green?** This is the fraction of the Broad_mask without pixels belonging to the Green_mask. It increases as more Broad_mask pixels leave the green class and therefore provides a readout of the green-to-non-green transition. |
| GLI — Green Leaf Index | $(2G - R - B) / (2G + R + B)$ | **How strongly does green dominate over red and blue within the green-classified pixels?** GLI quantifies relative greenness from the original RGB values of pixels selected by the Green_mask. Higher values indicate a stronger contribution of the green channel relative to red and blue. GLI describes colour only and does not depend on the size of the green area. |
| H_mean - Mean Hue of Green_mask pixels | $H\_mean = \Sigma H / \text{Green\_area}$ | **What is the average colour shade of the pixels in the Green_mask?**<br>H_mean describes the mean position of Green_mask pixels along the Hue axis and can therefore reveal a shift in colour, for example from green toward yellow-green. It describes colour only and does not depend on the size of the green area. |

**Table 2.** Experimental parameters and values to be finalized before submission. *Parameters compiled from the study’s laboratory records*

| Parameter | Status | Recovered value / note (source) |
| --- | --- | --- |
| Host / pathogen | Recovered | *A. thaliana* Col-0 (WT); *B. cinerea* B05.10 (nutrient/spore-number pathotests). |
| Conidial cryostock | Recovered | $1 \times 10^7$ conidia $\text{mL}^{-1}$ in 15% glycerol; $\sim 100\text{-}\mu\text{L}$ single-use aliquots at $-80^\circ\text{C}$ (*Botrytis* transformation lab book). |
| Conidial-harvest filtration | Recovered | 40-$\mu\text{m}$ nylon strainer (*Botrytis* transformation lab book). NB: one mémoire lists “0.4 $\mu\text{m}$”, almost certainly a typo for 40 $\mu\text{m}$. |
| Grouped-seedling density | Recovered | Standardized by volume: $\sim 10\text{ }\mu\text{L}$ dry seeds per $15 \times 15\text{ mm}$ nylon ($\approx 55$ seeds) (Ness; $\sim 11\text{ }\mu\text{L}$/nylon in the nutrient pathotest). |
| Growth temperature | Recovered | $\sim 21^\circ\text{C}$ (nutrient pathotest; working reference in method). |
| Relative humidity | Recovered | $\sim 40\%$ (target average; phytotron RH was variable). Reported as 40% (Ness). |
| Photoperiod | Recovered | 16 h light / 8 h dark, $21^\circ\text{C}$ (Ness). |
| Light intensity | Recovered | $\sim 70\text{ }\mu\text{mol m}^{-2}\text{ s}^{-1}$ ($\mu\text{E}$) (Ness; phytotron reference not available). |
| S1 sterilization solution | Recovered | Lab book M2 recipe: 10 mL bleach (1.5 g bleach in 40 mL water) + 40 mL absolute ethanol + 2 drops Tween 20; no SDS. |
| Inoculation & dilution | Recovered | Single 10-$\mu\text{L}$ central drop, $\sim 25$ conidia ($2.5 \times 10^3\text{ mL}^{-1}$). A method-development format ($5 \times 6.6\text{ }\mu\text{L}$ spots, $\sim 10^3\text{ mL}^{-1}$) delivered a similar low dose ($\sim 33$ conidia total); the single central drop was retained for throughput. |
| UV sporulation duration | To confirm | 2 d dark, then 9 d under 365-nm UV, at $21^\circ\text{C}$ (11 d total). |
| Post-freeze germination threshold | Recovered | $\geq 95\%$ (typically $\sim 100\%$); below this, use a fresher stock or re-sporulate (Ness). |
| Technical-validation panels | Decision pending | Decide which panels (JPEG/RAW, WB/device, manual segmentation, colour chart) go to the main paper vs supplementary. |

GreenDayZ – Phenotyping accepts JPEG, PNG, TIFF and DNG inputs and standardizes them to a common TIFF working format before downstream analysis. DataScan first performs dataset quality control before image preprocessing begins. Sample identifiers are parsed to associate each image with its experimental condition, replicate and acquisition day, allowing GreenDayz to reconstruct and summarize the complete experimental series. This step identifies missing time points or replicates, duplicate sample codes and suspicious filenames, while image readability and file integrity are checked in parallel. Problematic files can therefore be quarantined in a dedicated “compost” folder before batch processing, reducing the risk of interrupted or incomplete analyses.

Image preprocessing is organized into a set of independent image-processing modules, each implemented as a dedicated script and integrated into the GreenDayZ – Phenotyping graphical interface. This modular architecture allows individual processing steps to be tested, modified or extended independently, facilitating maintenance and future development of the application. The preprocessing workflow currently combines Convert & Crop, REFSET and White Balance. Convert & Crop standardizes image format and framing: RAW/DNG/JPEG acquisitions are converted to a common TIFF format when required, and the circular boundary of the Petri dish is automatically detected to recenter and crop each image to a standardized field of view. This reduces file size, limits framing differences between acquisitions and facilitates subsequent region-of-interest (ROI) detection. For large longitudinal datasets, REFSET provides a rapid preliminary assessment of acquisition variability before normalization is applied to the complete image series. A small representative subset, typically sampling across acquisition days, is extracted so that changes in ambient illumination, camera settings or imaging device can be evaluated without first processing hundreds of images. This inter-day sampling is particularly useful because such technical variation is most likely to occur between acquisition sessions. White Balance then corrects acquisition-related colour variation using one to four regions sampled from the white reference surrounding the Petri dish. For each REFSET image, GreenDayz measures the RGB values of these regions and derives channel-specific correction coefficients relative to a common neutral reference. The corrected REFSET is first inspected visually; once the correction is accepted, the same reference-based adjustment is applied to the corresponding cropped images in the complete series. This preprocessing step aims to reduce technical colour shifts before downstream biological colour segmentation and feature extraction.

### Image processing and feature extraction

GreenDayZ – Phenotyping colorimetry is based on the HSV (Hue, Saturation, Value/Brightness) colour space. Hue describes the dominant colour family, Saturation its chromatic purity relative to grey, and Value its brightness. These three dimensions are combined as threshold criteria to define binary pixel masks, which can be overlaid on the source image for visual validation before quantitative extraction.

Two independently parameterized masks were defined for the seedling assay. The Green_mask selects pixels corresponding to an operationally defined green phenotype, whereas the Broad_mask covers a broader colour-transition range encompassing pixels that progressively depart from this green phenotype. Because the masks are defined independently, they can overlap, particularly for intermediate yellow-green or khaki colours, commonly observed during disease progression. Their intersection is therefore retained as Overlap_area. Seed coats and some hypocotyl regions may also fall within the Broad_mask independently of infection, emphasizing that these masks represent operational colour classes rather than predefined biological tissue states. Raw images and mask overlays are consequently retained as quality-control outputs. GreenDayZ first extracts raw measurements from each image, including ROI area, Green_mask area (Green_area), Broad_mask area (Broad_area), their spatial intersection (Overlap_area), and mean RGB and HSV values within the Green_mask. These measurements are then used directly or combined to generate the core colorimetric readouts, including Green area fraction, Green-state fraction, Non-green fraction, GLI and H_mean (Table 1). Together, these readouts distinguish changes in the amount of green-classified tissue from changes in the colour properties of the remaining green pixels.

### Visual and analytical reporting

GreenDayz converts the extracted image features into visual and quantitative outputs suitable for longitudinal interpretation. Representative image boards and GIF animations provide a rapid overview of phenotype progression across the experiment, while the quantitative outputs can be plotted over time at the individual-sample or condition level. GreenDayz further supports longitudinal plotting and statistical comparison of selected metrics, while retaining access to individual-sample trajectories and the corresponding quality-control outputs.

### Application to the Arabidopsis–Botrytis pathosystem

Having established the GreenDayZ – Phenotyping workflow from dataset quality control to quantitative reporting, we applied it to the longitudinal A. thaliana Col-0–B. cinerea B05.10 seedling pathosystem to determine whether the selected image-derived readouts captured disease progression over time (Fig. 4). Prior to phenotypic analysis, inoculum quality was controlled by assessing post-freeze conidial germination and by quantifying the number of conidia effectively delivered per inoculation drop.

**Figure 4.**
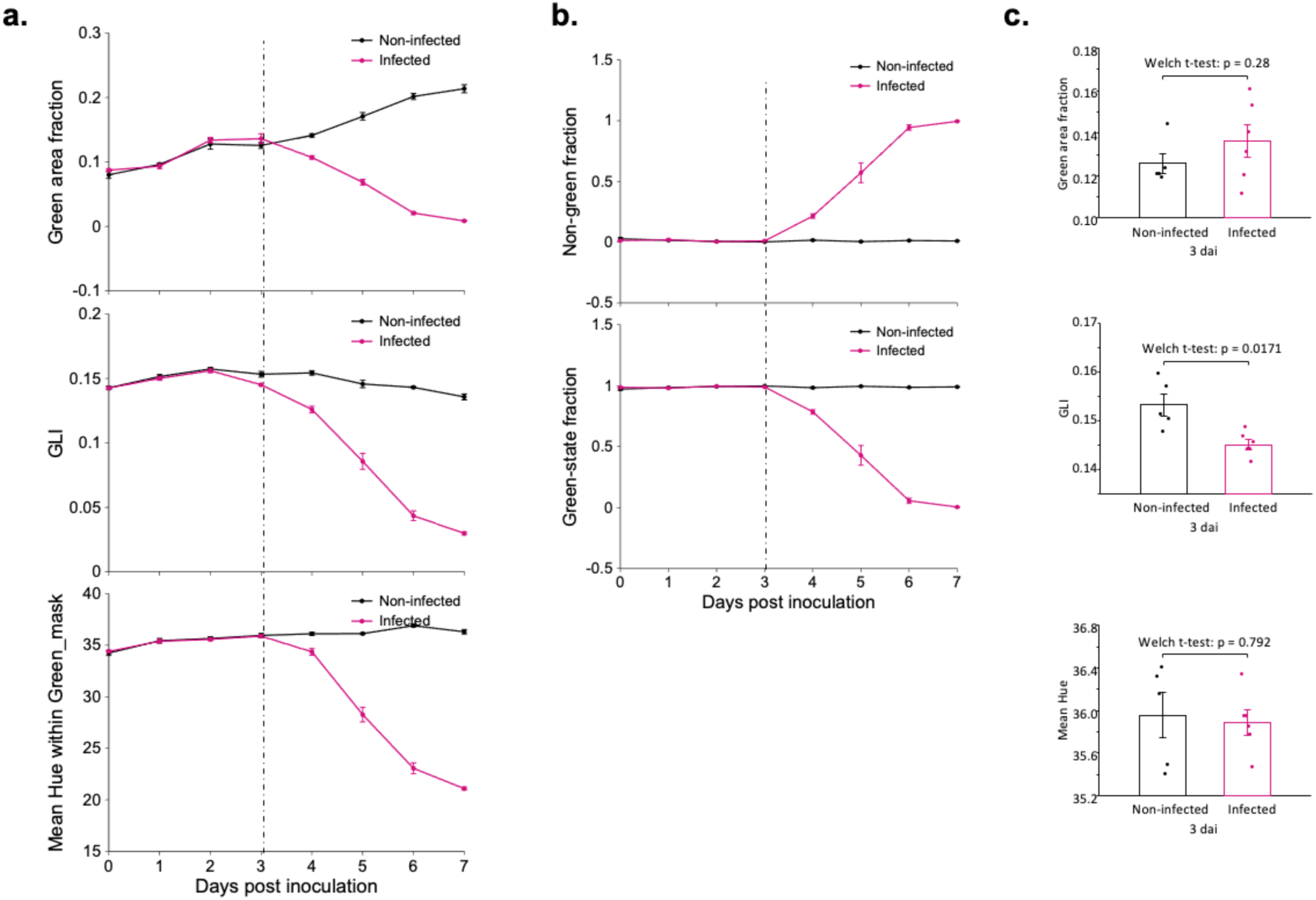
Data Analysis outputs reveal longitudinal divergence between non-infected and infected seedlings, with only the colour-based readout already altered at 3 dai. GreenDayZ Data Analysis outputs comparing non-infected and *Botrytis cinerea*-infected *Arabidopsis thaliana* seedlings over the infection time course (n = 6 biological replicates per condition). (a) Longitudinal evolution of Green area fraction, GLI and mean Hue within the Green_mask. The three readouts are superimposable until 3 dai, then diverge progressively as infection develops. (b) Longitudinal evolution of the Non-green fraction and the Green-state fraction, which capture the same transition as a shift of the seedling population between colour classes. (c) The three readouts of panel a compared at 3 dai, before visible divergence. Green area fraction and mean Hue remained similar between conditions, whereas GLI showed a modest but detectable difference, indicating that a change in the relative greenness of the remaining green-classified tissue can be detected before a reduction in green area or a shift in mean hue becomes apparent. Dashed vertical lines mark 3 dai. Points and bars are means ± SEM, dots are individual biological replicates, and p-values are from two-sided Welch t-tests.

Across the time course, non-infected seedlings showed an increasing Green area fraction *(fraction of ROI pixels belonging to the Green_mask)* and comparatively stable GLI *(relative dominance of the G channel over R and B within Green_mask pixels)* and H_mean *(mean Hue within the Green_mask pixels)*, whereas infected seedlings progressively diverged after the first days of interaction (Fig. 4a). In infected seedlings, Green area fraction decreased to very low values by 6–7 dai, GLI declined in parallel, while H_mean remained initially stable before shifting at later time points. The green-to-non-green transition was also captured by the Green-state fraction and Non-green fraction (Fig. 4b). Green-state fraction *(fraction of Broad_mask pixels also belonging to the Green_mask)* remained high early in the time course and then decreased sharply in infected seedlings. Conversely, Non-green fraction *(fraction of Broad_mask pixels not belonging to the Green_mask)* increased toward 1. Non-infected controls remained comparatively stable. Because the two indices are complementary, they describe the same transition in opposite directions. To examine whether an earlier colorimetric difference could be detected before these marked longitudinal divergences became apparent, Green area fraction, GLI and H_mean were compared specifically at 3 dai (Fig. 4c). Green area fraction did not differ significantly between non-infected and infected seedlings (Welch’s *t*-test, *P* = 0.280), and no difference was detected for H_mean (*P* = 0.792). In contrast, GLI was already significantly lower in infected seedlings (*P* = 0.0171). Thus, at this early stage, the relative RGB greenness of Green_mask pixels was altered before either a measurable loss of projected green area or a detectable shift in mean Hue. This early decrease in GLI may therefore reflect subtle changes in pigment balance within tissues that still appear macroscopically green, rather than an overt loss of green tissue. Together, these results show that the selected GreenDayZ readouts capture complementary dimensions of disease progression, from early changes in pixel greenness to the later loss of green-classified tissue.

Large longitudinal image datasets can rapidly become storage-intensive, particularly when high-resolution TIFF files are retained throughout the analysis workflow. Image-format robustness was therefore assessed independently from the biological comparison. The infected time series was re-analysed after conversion of the reference TIFF images to smartphone-like JPEG files using a quality setting of 90 (Supplementary Fig. S3). JPEG compression reduced the size of a representative image from approximately 57-fold reduction in file size (from 22.8 MB to 0.40 MB), while Green area fraction, GLI, Green-state fraction and Non-green fraction remained nearly superimposable between formats throughout the time course. Under the tested compression conditions, the large reduction in file size therefore had little effect on the selected GreenDayZ - Phenotyping readouts. For a dataset of 84 images, representative of the scale of the experiment presented here, this corresponds to an approximate reduction from 1.9 GB of TIFF data to only 34 MB of JPEG files.

## Discussion

This work combines a miniaturized, low-inoculum *Arabidopsis thaliana - Botrytis cinerea* seedling pathosystem, a standardized smartphone imaging setup and GreenDayZ - Phenotyping for longitudinal RGB analysis. Together, these components enable non-destructive monitoring of disease progression while preserving parallel biological material for destructive molecular analyses. The approach is particularly suited to kinetic experiments in which phenotypic trajectories need to be related to independent measurements of fungal growth or host responses.

The use of young seedlings together with a very low inoculum provides access to an experimental window that remains less explored than infection of mature leaves under higher pathogen loads. In this context, GreenDayZ captures complementary aspects of disease progression, including changes in projected green area, RGB greenness, Hue and the transition from green to non-green colour classes. These trajectories can be aligned with independent measurements such as bioluminescence, qPCR, phytohormone profiling or transcriptomics, providing a framework to relate phenotypic changes to the early stages of infection. The biological system also has several specific features that define its scale and experimental context. Grouped seedlings provide a population-level rather than single-plant readout, while the central inoculation site imposes a defined spatial origin of infection. The grouped configuration may also generate a particular local microenvironment around the seedlings; accordingly, relative humidity in the growth chamber should be carefully controlled and kept consistent between experiments, as humidity strongly influences plant–pathogen interactions and disease development. Depending on the biological question, seedling organization or inoculation geometry could be adapted without changing the overall imaging and analysis framework.

The imaging workflow was intentionally kept simple and accessible. Standardized positioning, controlled illumination and white-reference normalization reduce acquisition variability, while visual inspection is retained throughout the analysis. The close agreement between TIFF- and JPEG-derived trajectories further indicates that standard JPEG acquisition is sufficient for the readouts tested here, despite a substantial reduction in file size. This is particularly relevant for large longitudinal datasets and supports routine smartphone-based acquisition. GreenDayZ - Phenotyping should therefore be considered a functional but not yet finalized implementation. The core modules required for dataset quality control, image preprocessing, colorimetric segmentation and longitudinal analysis are operational, whereas additional functions, including full colour-chart calibration and automated sample identification, remain under development. Sample identification is currently performed manually using the code visible in each image; a QR-code-based system is being developed as a more robust alternative for automated tracking.

Overall, this framework provides a practical means of linking low-inoculum seedling infection with longitudinal phenotyping and molecular sampling, and should facilitate the study of early stages of pathogen infection before extensive symptom development.

## Methods

### Plant material, medium and seed sterilization

*Arabidopsis thaliana* (Col-0) seeds were grown on sucrose-free Murashige and Skoog medium (Duchefa Biochemie #M0231.005) (9), supplemented with Gamborg B5 vitamins (2.2 g L⁻¹) and plant agar (0.9%, w/v), adjusted to pH 5.7 with KOH. Exactly 8 mL of medium was dispensed into each 55-mm Petri dish; closed promptly and stored at 4°C until use. Seeds were surface-sterilized in 50-mL tubes using freshly prepared S1 solution [10 mL bleach (1.5 g bleach dissolved in 40 mL water) plus 40 mL absolute ethanol and 2 drops of Tween 20]. Up to 1 mL dry seeds was processed per tube (falcon 50mL) to maintain an adequate seed-to-solution ratio. Seeds were gently inverted in S1 for 5 min, washed twice in 70% ethanol for 5 min, transferred to approximately 15 mL absolute ethanol, recovered using a widened sterile P1000 tip and dried for approximately 5 min between two sterile Whatman layers. Dry, free-flowing seeds were used for sowing.

### Nylon-supported sowing and seedling establishment

For grouped assays, seeds were distributed onto autoclaved 100µm nylon squares (15 × 15 mm). During sowing, several nylon squares were arranged in a sterile square Petri dish (120 x 120 x 17 mm) and temporarily covered with a sterilized 1-mm sowing grid made from mosquito-net mesh (Appendix 1). Dry seeds were sprinkled using a simple Falcon-tube seed shaker fitted with sterile gauze (4 layers). Seed input was standardized by volume at approximately 10 µL of dry seeds per 15 × 15 mm nylon square (corresponding to ∼55 seeds). The grid was removed in a single continuous motion to limit local aggregation caused by the water film forming at the humid nylon/agar interface. Seeded nylon squares were then transferred individually on 55mm MSk plates. Initial seed density and spatial distribution were visually inspected; when dispersion was not homogeneous, extreme plates were excluded where necessary and remaining dense or sparse plates were distributed across experimental conditions and replicate groups. Seeded plates were stratified for 2-3 days at 4°C in darkness, exposed to light for 2-6 h at approximately 21°C to initiate germination, and then returned to light until cotyledons were fully green (typically 1-2 days). Because hypocotyl elongation is strongly temperature-dependent and accelerates at 24°C, and because excessive dark growth promotes lodging and cotyledon-reserve depletion, temperature and duration were carefully checked and kept constant within each experiment. Seed germination and developmental synchrony were visually checked during establishment; plates showing contamination, strongly abnormal germination or marked developmental heterogeneity were excluded.

### Botrytis cinerea culture, conidial stock preparation and quality control

*Botrytis cinerea* (strain B05.10) **[10]** was pre-cultured for 2-3 days in darkness at 21°C. A plug was excised from the actively growing colony margin and transferred to malt sporulation (MS) medium containing glucose (5 g L⁻¹), malt extract (20 g L⁻¹), bacto tryptone (1 g L⁻¹), casamino acids (1 g L⁻¹), yeast extract (1 g L⁻¹), RNA extract (0.2 g L⁻¹) and agar (16 g L⁻¹), pH 5. Sporulation plates were incubated in darkness (2 days ; 21°C) followed by 365-nm UV exposition (9 days ; 21°C). Conidia were recovered from the colony surface in 2-3 mL sterile autoclaved water using a sterile plastic spreader and filtered (40-µm nylon strainer) to remove mycelial fragments. Conidia were counted using a Thoma chamber, adjusted to approximately 1 × 10⁷ conidia mL⁻¹ in 15% glycerol, distributed into ∼100 µL single-use aliquots and stored at -80°C. Thawed aliquots were not refrozen. Each newly prepared frozen stock was subjected to a post-freeze germination quality control: a dedicated aliquot was thawed, and 10 µL was deposited and distributed on MS agar to generate countable germination events/microcolonies, assessed after 1-2 days and recorded with the stock batch identifier. Stocks were accepted when post-freeze conidial germination was ≥ 95% (typically ∼100%); below this threshold, a fresher stock was used or a new stock was produced.

### Low-dose inoculation and pathotest conditions

On the day of inoculation, one single-use reference aliquot was thawed and serially diluted in sterile autoclaved water to a nominal concentration of 2.5 × 10³ conidia mL⁻¹, corresponding to approximately 25 conidia per 10 µL inoculum. The working suspension was maintained under constant magnetic stirring in a sterile flat-bottom vessel throughout inoculation to limit conidial sedimentation. A single 10 µL droplet delivering approximately 25 conidia was deposited at the centre of each nylon-supported seedling population; mock controls received sterile water. To estimate the actual delivered dose, 20 independent 10 µL aliquots were sampled from the same continuously stirred suspension and the conidia counted; this dose quality control was recorded for each inoculation session.

Inoculated plates were immediately returned to controlled growth conditions and maintained in open **crystallizing dishes** to reduce local airflow and humidity heterogeneity. Growth was carried out at approximately 21°C under a 16h light / 8h dark photoperiod at approximately 40% relative humidity, with a light intensity of approximately 70 µmol m⁻² s⁻¹.

### DIY photo station and longitudinal image acquisition

RGB images were acquired under the sterile microbiology hood using a consumer smartphone mounted above a simple DIY station. The plate support consisted of black and white Plexiglas layers that reproducibly positioned the 55 mm dish while retaining a visible white neutral-reference area; a separate black Plexiglas plate constrained smartphone position. Illumination was provided by a consumer LED ring light. Unnecessary ambient-light sources were switched off, smartphone flash was disabled and the ring-light mode and intensity were held constant within an experiment. A Bluetooth shutter was used to avoid touching the acquisition setup. Immediately before imaging, the Petri-dish lid was removed to reduce reflections; the plate was centred on the support and the coded lid was positioned above the plate so that sample identity remained visible in the image field without obscuring the seedlings. Original capture metadata were preserved; when supported by the device, timestamp-based filenames (e.g., YYYYMMDD_HHMMSS) were used to facilitate chronological traceability. RAW/DNG images were optionally acquired for technical validation or advanced colour-processing tests. Standard JPEG images were sufficient for routine analysis.

## Software reproducibility and data traceability

The GreenDayz version, experiment parameter preset and analysis metadata were archived with the quantitative outputs. The reproducibility package will include the frozen application release, a Python environment/dependency file, an installation guide, the parameter preset, a small example dataset and user instructions. Routine use of the graphical interface does not require programming after installation. xx

## CRedit authorship contribution statement

Conceptualization : F.-X.G ; Supervision : F.-X.G ; Investigation : I.T, F.-X.G. ; Methodology : I.T, F.-X.G. ; Visualization : I.T, F.-X.G. ; Data curation : F.-X.G. ; Software : F.-X.G. ; Writing – original draft : I.T, F.-X.G. ; Writing - review and editing : I.T, C.Br., N.T., M.C., C.Bo, F.-X.G.

---

**Figure S1.**
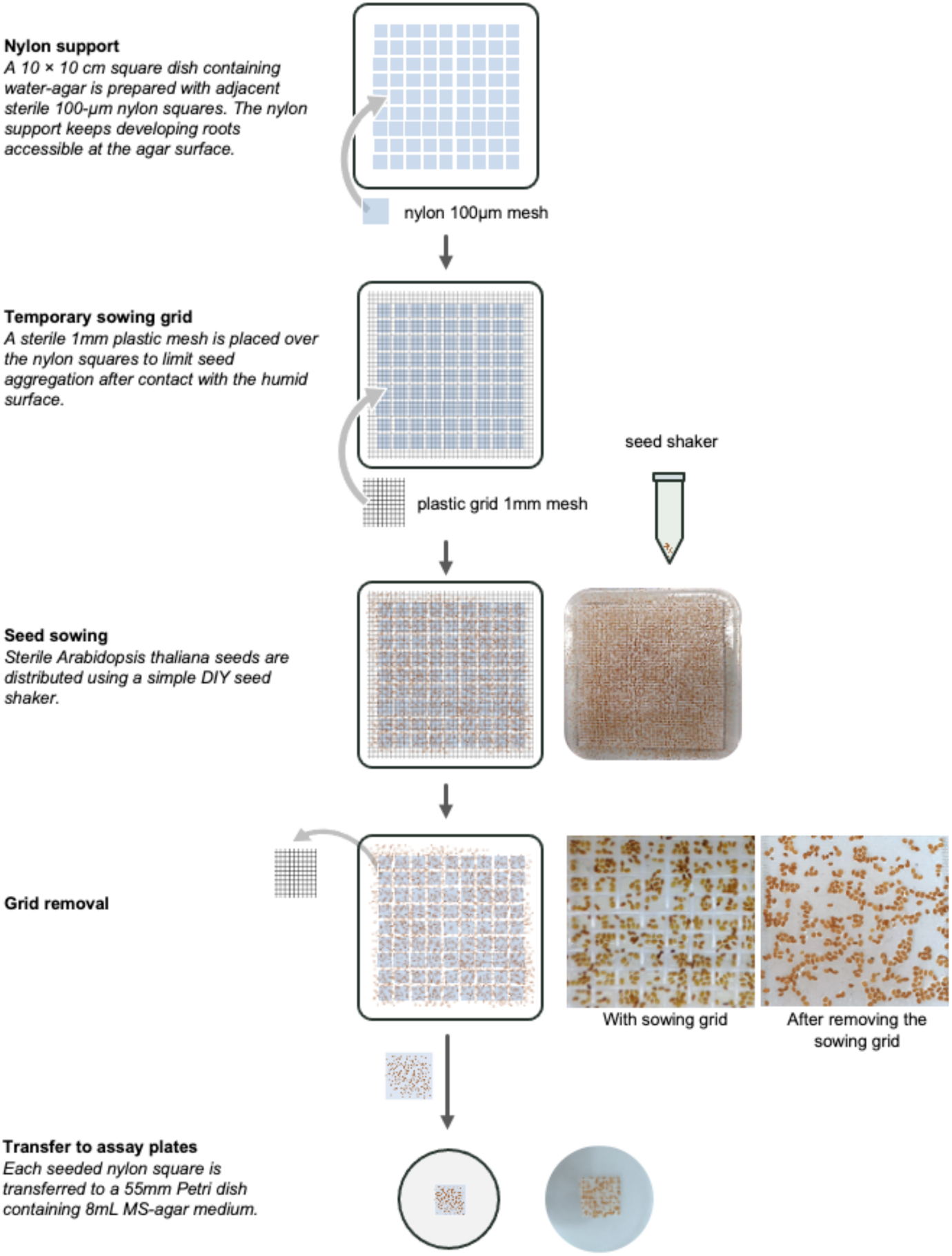
Nylon-supported seed sowing and transfer workflow. Sterile 100 µm nylon squares are arranged on water-agar in a 10 × 10 cm square dish. A temporary 1 mm sowing grid is placed over the nylon supports to limit seed aggregation during deposition. Sterile *Arabidopsis thaliana* seeds are distributed using a DIY seed shaker, after which the grid is removed and each seeded nylon square is transferred to the centre of a 55 mm Petri dish containing 8 mL MS-agar medium. Representative close-up images illustrate seed distribution before and after grid removal.

**Figure S2.**
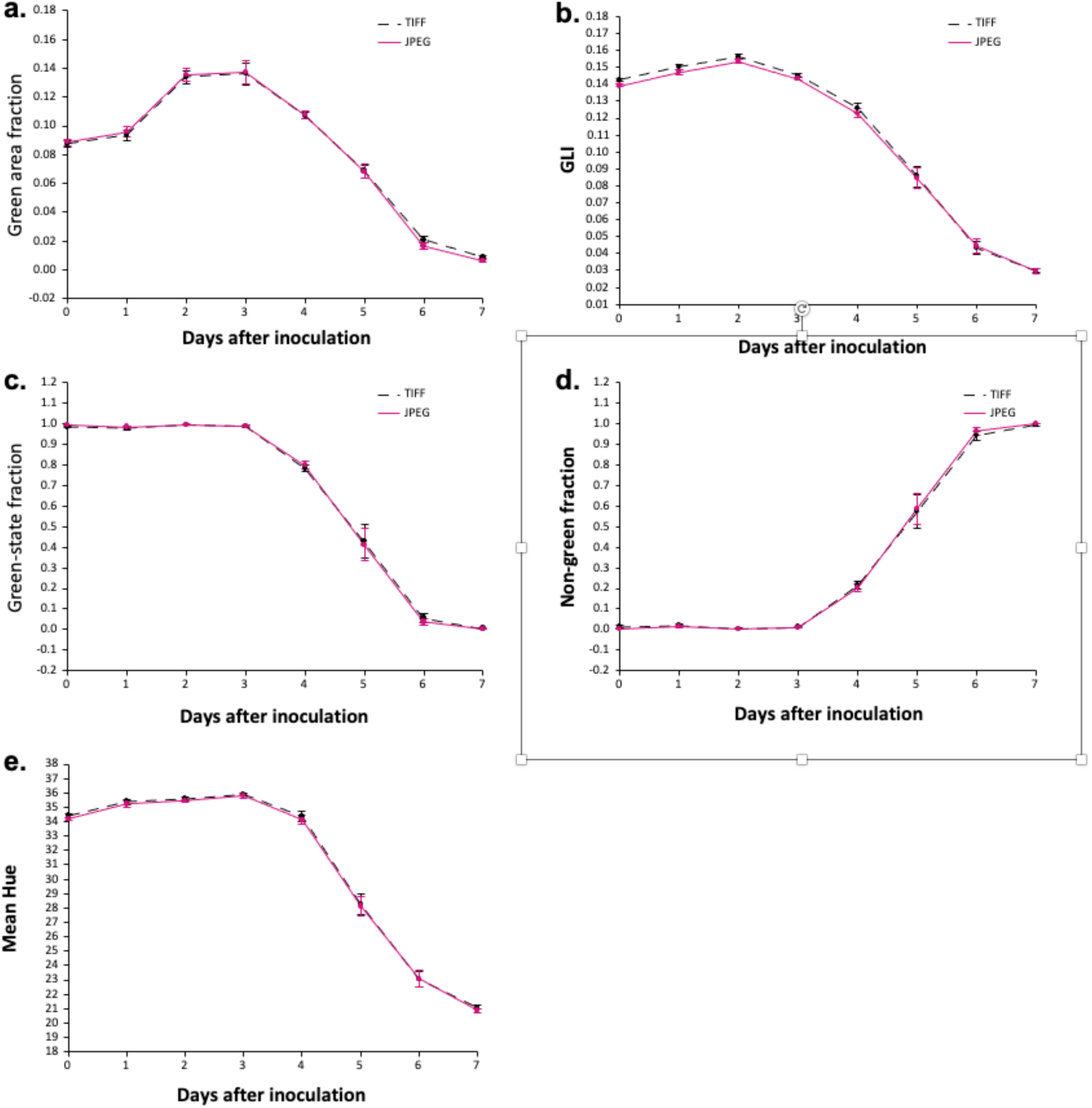
Effect of smartphone JPEG compression on longitudinal readouts. The same infected *Arabidopsis thaliana-Botrytis cinerea* image series was analysed from reference TIFF images (black dashed lines) and from JPEG-compressed versions of the same images (magenta solid lines). JPEG files were generated from the reference TIFF images using a quality setting of 90 and subsequently converted back to the common TIFF working format before application of the same Colorimetrics workflow. **(a)** Green area fraction, corresponding to the fraction of ROI pixels belonging to the Green_mask. **(b)** Green Leaf Index (GLI), reporting the relative dominance of the G channel over R and B within Green_mask pixels. **(c)** Green-state fraction, corresponding to the fraction of Broad_mask pixels also belonging to the Green_mask. **(d)** Non-green fraction, corresponding to the fraction of Broad_mask pixels not belonging to the Green_mask. **(e)** H_mean, representing the mean Hue within Green_mask pixels. Measurements are shown for infected seedlings from 0 to 7 days after inoculation (dai). Points represent means ± SEM (*n* = 6 biological replicates per time point). TIFF- and JPEG-derived trajectories were nearly superimposable for all five readouts, indicating that JPEG compression at the tested quality setting substantially reduced image size without materially altering the longitudinal GreenDayZ - Phenotyping measurements.

